# Bio-electric potentials in superior plants: electric collective behaviour

**DOI:** 10.1101/2022.04.07.487233

**Authors:** Alessandro Chiolerio, Mohammad Mahdi Dehshibi, Giuseppe Vitiello, Alessandro Bernard, Paolo Ceretto, Massimo Arvat, Andrew Adamatzky

## Abstract

Electrical activity is used by plants in long term signalling and information transfer between the distant parts of the plant. Biopotential recordings from trees in a natural environment have been so far less discussed in scientific literature. Here we present our data about the open science experiment TRee-hUMAn iNterface (TRUMAN) located in Paneveggio forest (Valle di Fiemme, Trento, Italy), cultivated since one thousand years for the production of harmonic wood from *Picea abies* (red fir). We show that: i) biopotential features based on xylem can be correlated with the solar (and lunar) cycle ii) dead tree logs show an electrical activity that is correlated with that of neighbouring trees iii) statistical features of the spike-like peaks are evidenced, including amplitude, frequency, propagation speed, entropy iv) a quantum field theory is presented to highlight the collective behaviour of the forest, supported by preliminar correlation analyses between electrical signal Kolmogorov entropy and thermographies Shannon entropy.

## 1. Introduction

The increasing demand for tree and forest health monitoring due to ongoing climate change requires new future-oriented and nondestructive measurement techniques [1, 2, 3, 4, 5]. One such example is given by Electrical Resistivity Tomography (ERT), providing insights into living trees based on resistivity measurements performed with cross-sectional distribution [6, 7, 8]. External factors, such as temperature, and water status, have been evaluated also in *Picea abies* individuals [9]. Trees are renowned for their salient intelligence, capabilities to implement distributed information processing, showing indicators of advanced perception, cognition and adaptive behaviour, anticipatory responses and swarm intelligence [10, 11, 12, 13, 14]. Trees employ impulses of electrical activity to coordinate actions of their bodies and long-distance communication [11, 15, 16]. The bursts of impulses could be either endoge-nous, e.g. related to motor activities or in a response to external stimulation, e.g. temperature, osmotic environment, rain, wind, mechanical stimulation. Electric signals could propagate between any types of cells in a plant tissue, there are indications, however, of higher conductivity of the vascular system which might act as a network of pathways for travelling electrical impulses. We can speculate that when several impulses travel along a vascular network they meet and interact with other impulses: for example in a tree trunk the vertical impulse movement is triggered by cytoplasm migration, while the horizontal cross-talk is granted by ion diffusion.

With the aim of developing a living information processing system, we monitor the electric potential expressed by the plants in a finite number of sites, to observe the entire forest and not a single plant. The electrical response is dependent on the state of the ecosystem that the root structure is embedded into, particularly to the water hydrostatic pressure in the soil. By analysing the response, one can infer about the connection graph and the features of environmental and health conditions. Moreover, details about collective behaviour of the plants can also be highlighted. To implement such living computing network we have selected a portion of Valle di Fiemme forest located in Paneveggio (TN, Italy) [17, 18, 19, 20] including five spots with trees in proximity, of a proper age, spanning approximately 8000 square meters (see Figure 1). Such forest-based information processing devices can be few hundreds of kilometres in diameter and used to analyse and monitor huge underground ecosystems in an environment friendly way. It could be used as a universal query machine to infer morphological features of the root system in real-time: e.g. its shape, depth, connectivity patterns, typical root size, distribution in space and other, or to query about the health state of the ecosystem.

**Figure 1.**
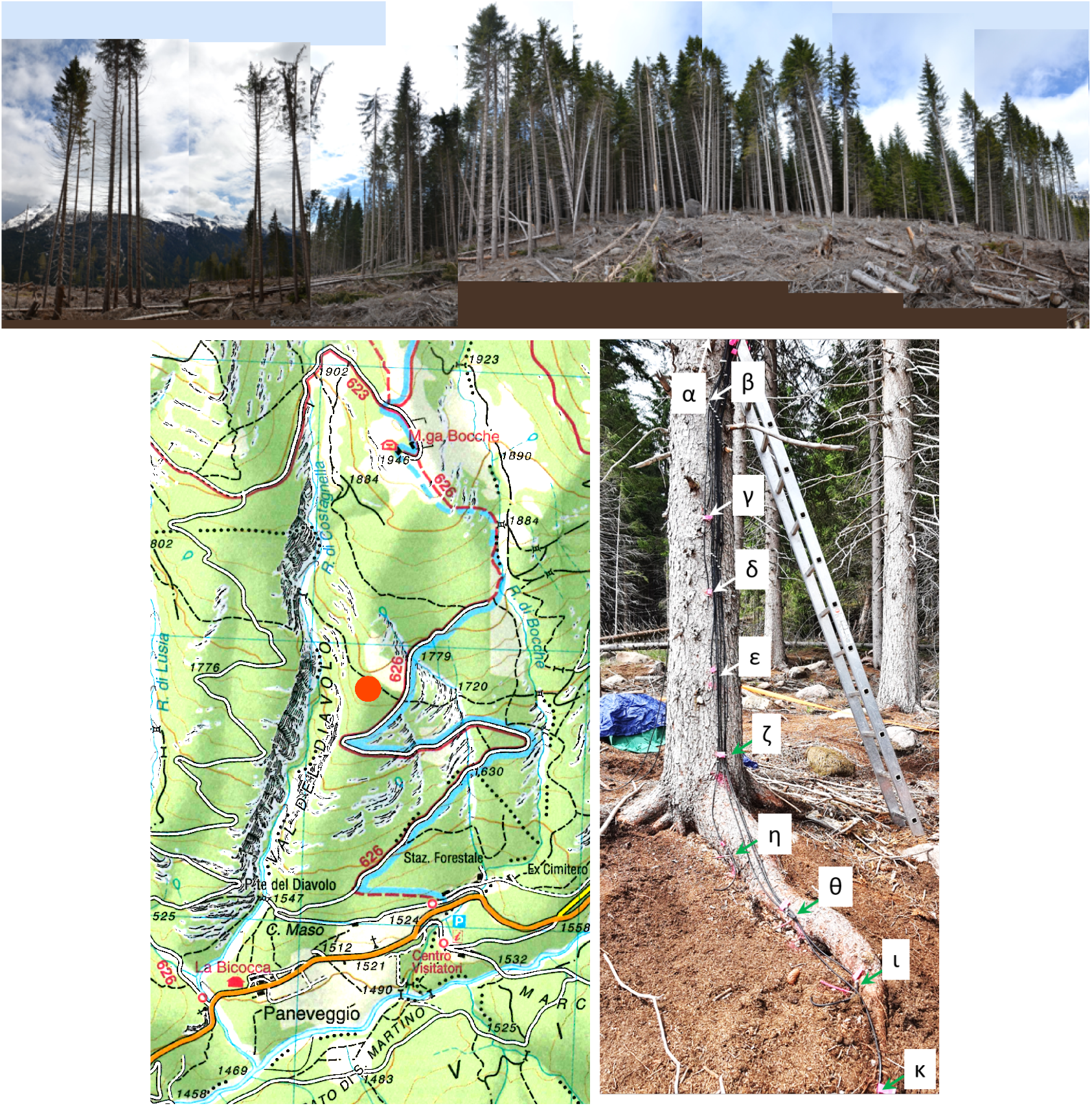
Top image: area of the installation in Paneveggio (TN, Italy) seen with a patchwork of camera shots. Bottom, from left to right: map showing the exact location of the experiment, indicated by the red circle (diameter corresponding to 100 m); detail showing the geometry of the electrodes inserted in a *Picea abies* tree of approximately 70 years of age, labelled with greek letters; detail about the box hosting PV modules, battery for powering the infrastructure and WiFi router for collection and data relay, during installation.

The paper is organised as follows. Experimental setup is presented in Sect. 2. Electrical activity and electrical properties of trees are analysed in Sect. 3. A theoretical modelling of the forest as a collective organism is sketched on the basis of a quantum field theory framework in Sect. 4. We draw the conclusions in Sect. 5.

## 2. Materials and methods

### 2.1. Experimental

Bio-potentials were collected using stainless steel (AISI 316) threaded rods of 6 mm diameter, positioned along the trunk, uncovered portion of roots and logs, evenly spaced by 50 cm. The length of the bores was chosen to allow a perfect vertical alignment between the tips of the electrodes. They were isolated, keeping uncovered and electrically conductive the probing tip (1 cm in length) and the back (2 cm in length) where soldering cable lugs were bolted. Each electrode was connected using double shielded ultra low resistance cable INCA1050HPLC from MD Italy for high fidelity audio application, positioned along the trunk using fairleds. A DI-710-US data logger was used in individual channel mode or differential mode, recording at a frequency of 10 Samples/s per channel, with 16 bit resolution and 100 mV of voltage range. Data were smoothed using a Savitzky-Golay first order function over a variable size window, ranging between 41 and 301 points, depending on the noise floor. Electrodes were labelled with Greek letters:*α, β, γ, δ, ϵ, η, ζ, θ, ι, κ*. Collection sites were labelled with Latin letters: A, B, C, D, and E. Trees belonging to a collection site were labelled with Arabic numbers: 1, 2, 3 and 4. Resins electronic properties were measured using a A Keithley 2635A multimeter for DC characterization (IV curves in the range ±200 mV and cycles in the range ±200 V) and an Agilent E4980A precision LCR meter for AC characterization (range from 20 Hz up to 2 MHz, 1 *V*_*RMS*_).

Electrical activity is recorded using a number of electrodes in mechanical connection with the xilematic tissue, regularly spaced along the trunks, where possible extending down to the roots of the trees, and eventually also to the dead logs in the surroundings. Each site hosts 10 electrodes connected to a data logger to collect information in differential mode. Typical measurement chunks cover approximately the duration of a day and highlight the features of superior plant potential oscillations (see Figure 2): the typical bias level varies between 5 and 150 mV, the fluctuation spike-like peak occurs with temporal spacing of 15 ± 50 s (monomodal distribution, median: 4 s, skewness 13, kurtosis 300, maximum interval 1600 s) and has an average amplitude of 10 ± 8 mV (monomodal distribution, mode: 1.5 mV, skewness 2.1, kurtosis 5.8, maximum amplitude 46 mV). Broader structures, lasting tens of minutes, can be associated to the occurrence of sunrise. The diurnal features of biosignal are clearly distinguishable from the nocturnal ones, including a different predominance of low and high frequency noises, as well as a higher number of spikes during night. By tracing sharp peaks it is possible to observe the propagation delay of stimuli, evidencing the descending vascular path probed by the electrodes.

**Figure 2.**
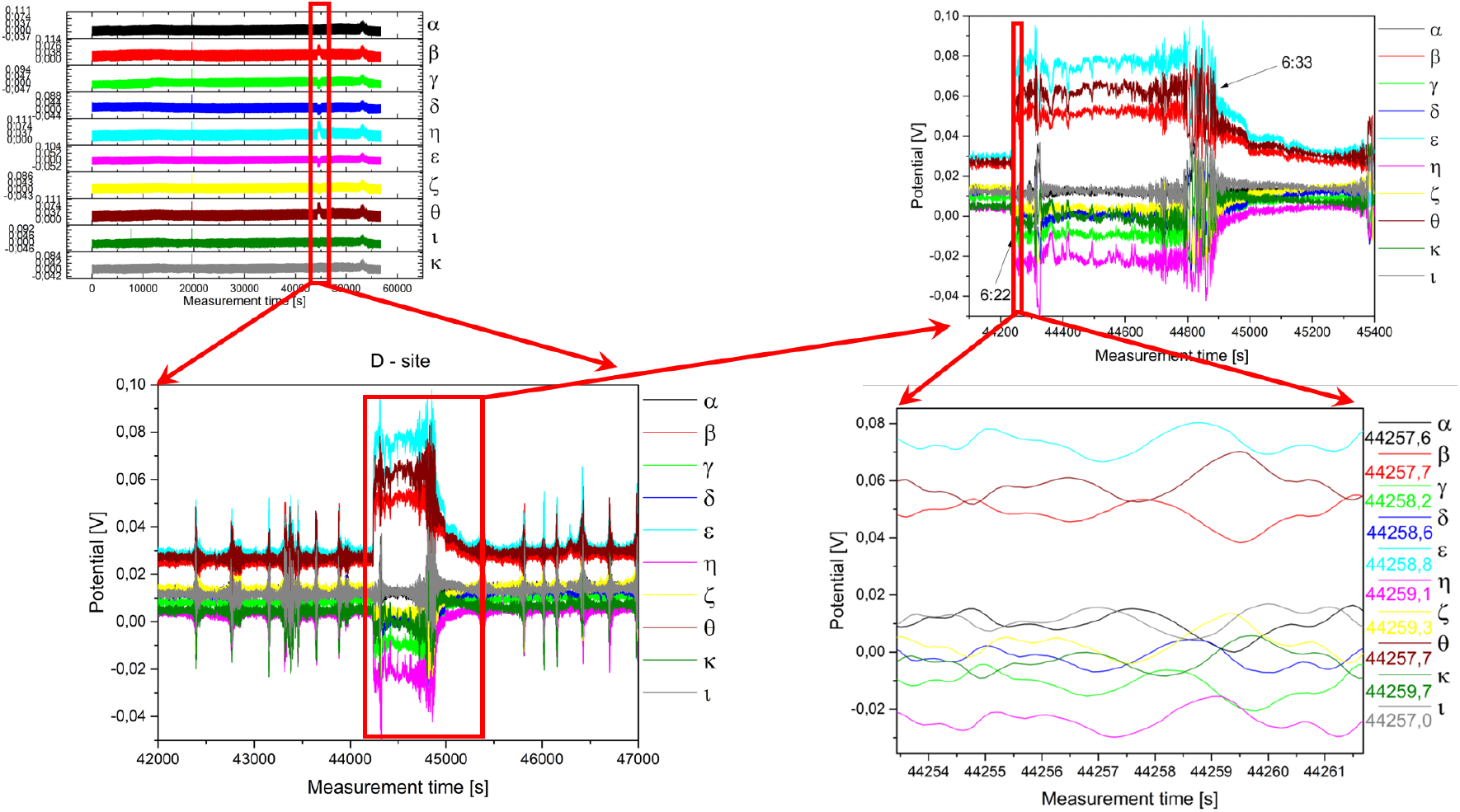
Data measured from site D, tree D1, over ten individual electrodes. Four magnification steps are shown.

### 2.2. Analysis

We formulated our problem as a Bayesian probability because we are dealing with inferential statistics since we do not have precise information about the underlying distribution of the electrical responses collected from our sensory setup (*i*.*e*., unobserved variable). The cumulative distribution function (CDF) depicts the probability that a random variable *X* can be found with a value equal to or less than a particular The CDF’s inverse distribution function, on the other hand, expresses the reciprocal of a random variable and provides the value associated with a certain cumulative probability. We utilise it to find spike occurrences in the Bayesian context of prior distributions and posterior distributions for scale parameters because we are dealing with modelling phenomena where numerically significant values are more probable than the case for the normal distribution.

For the acquired signal *x*, we use the Eq. 1 to calculate the inverse of the standard normal CDF.

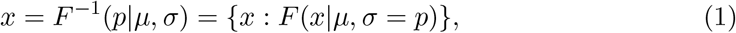

where *µ* is the mean, *σ* is the standard deviation, and 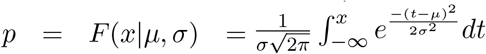. The standard normal distribution has zero mean and unit stan-dard deviation, and the result *x* is the solution of the integral equation where we supply the desired probability *p*.

Assume 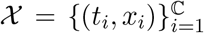 is a recording set of C channels with the entire length of *T* seconds and samplings rate of *f*_*s*_ Hz, where *x* = max(*x*) − *x* defines the signal’s sample value at time *t*, 1 ≤ *t* ≤ *T*. Our objective is to detect the set of spike events 𝒮 = {*s*_1_, *s*_2_, …, *s*_*η*_}, where *η << T*. A spike event (*i*.*e*., a local peak) is a data sample that is either greater than the two neighbouring samples or equal to ∞. We calculate the following complexity measurements by locating the local peaks.

1. Spikes number: Total number of located peaks.
2. Barcode entropy: We represent the entire recording duration *T* with a binary string 𝒮, with ‘1s’ indicating the availability of spike events and ‘0s’ otherwise. The Shannon entropy of this binary string (i.e., barcode entropy) is calculated by *H*(𝒮) = − ∑_*i*_ *p*(*s*_*i*_) log *p*(*s*_*i*_).
3. Simpson index: It is calculated as *Simpson* = ∑_*w* ∈*W*_ (*ν*(*w*)*/η*)^2^. It linearly correlates with Shannon entropy for *H <* 3 and the relationship becomes logarithmic for higher values of *H*. The value of *Simpson* ranges between 0 and 1, where 1 represents infinite diversity and 0, no diversity.
4. Space filling (*D*) is the ratio of non-zero entries in *W* to the total length of string.
5. Expressiveness (*E*) is calculated as the Shannon entropy *H* divided by space-filling ratio *D*, where it reflects the ‘economy of diversity’.
6. Lempel–Ziv complexity (*LZ*) is used to assess temporal signal diversity, i.e., compressibility. We use Kolmogorov complexity algorithm [21] to measure the Lempel–Ziv complexity.
7. Rényi (*R*_*q*_) [22] and Tsallis (*Tq*) [23] additive entropy concepts are generalisations of the classical Shannon entropy. Regardless of the generalisation, these two entropy measurements are used in conjunction with the Principle of maximum entropy, with entropy’s main application being in statistical estimation theory. Tsallis and Rényi entropy measurements are expressed as 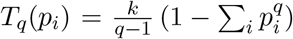 and 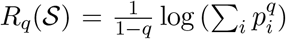, where *q* is the *entropic-index* or Rényi entropy order with *q* ≥ 0 and *q* ≠ 1, which in our experiments was set to 2.

## 3. Results

When measurements are studied in the frequency domain by taking the Fast Fourier Transform (FFT), the outcomes are that above approximately 200 mHz the Total Impulse Square Amplitude becomes almost perfectly horizontal and shows no clear features, while at lower frequencies typical features appear showing fingerprints in the 1 to 10 mHz range (see Figure 3). The Short Time Fourier Transform (STFT) is also a powerful tool to visualize the different features of diurnal signals versus nocturnal ones. The different pace, showing a high frequency nocturnal activity and a more quiet diurnal one, can be justified by a higher degree of inner disorder and will be discussed in Sect. 4.

**Figure 3.**
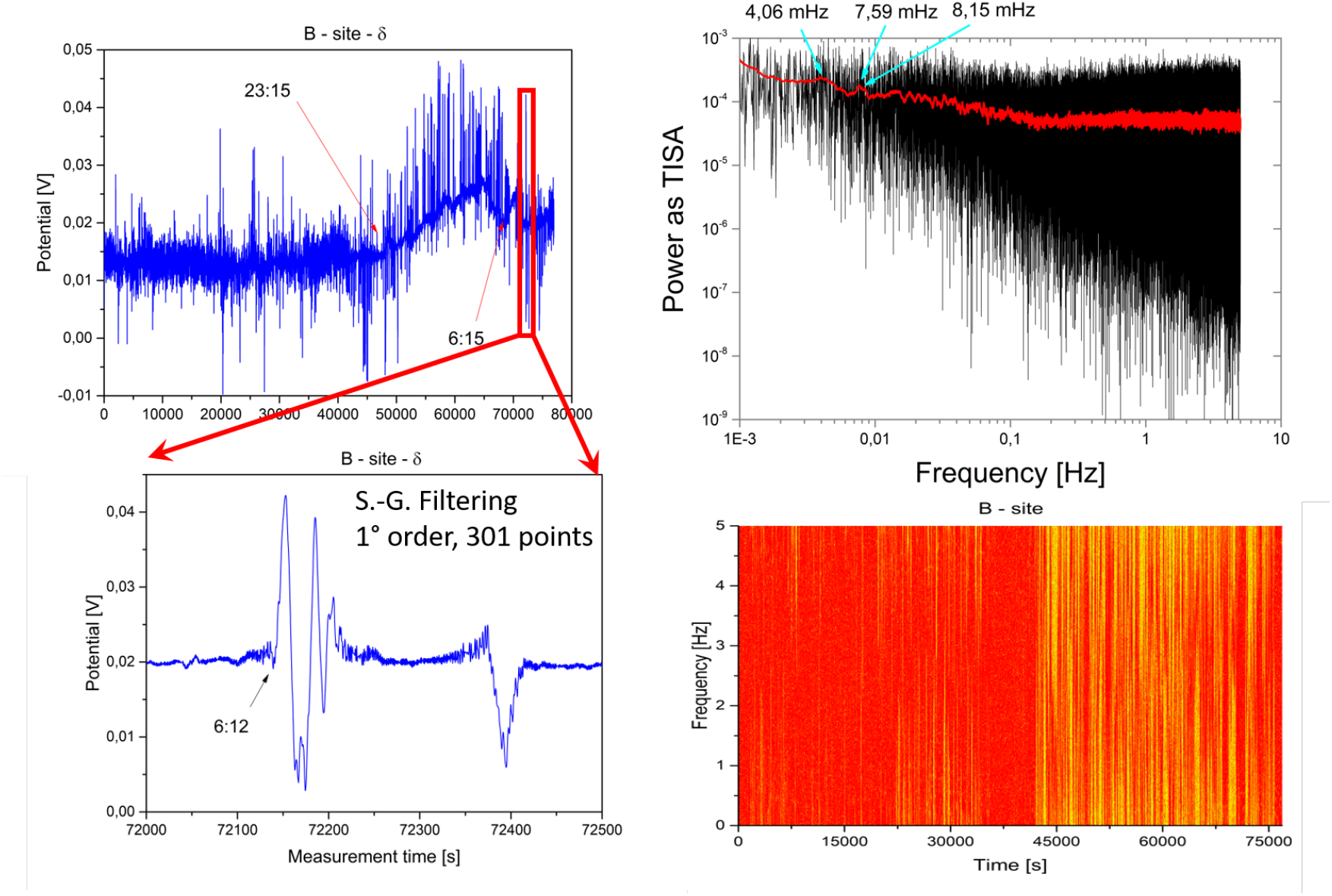
Data measured from site B, tree B2, over ten individual electrodes. Two magnification steps are shown, including the Fast Fourier Transform analysis as Total Impulse Square Amplitude and the Short Time Fourier Transform map.

The effects of sun and moon movement can also be sensed, in particular Figure 4 reports the stacked curves and some magnified portions of one single channel, to evidence the differences in noise and spike patterns of the signal. Unfortunately sunset / moon rise and sunrise / moon set are too close to clearly resolve in time their effects on the electrical response. The propagation speed of the pseudo peaks along the channels has been calculated to be 0.125 ± 0.005 m/s.

**Figure 4.**
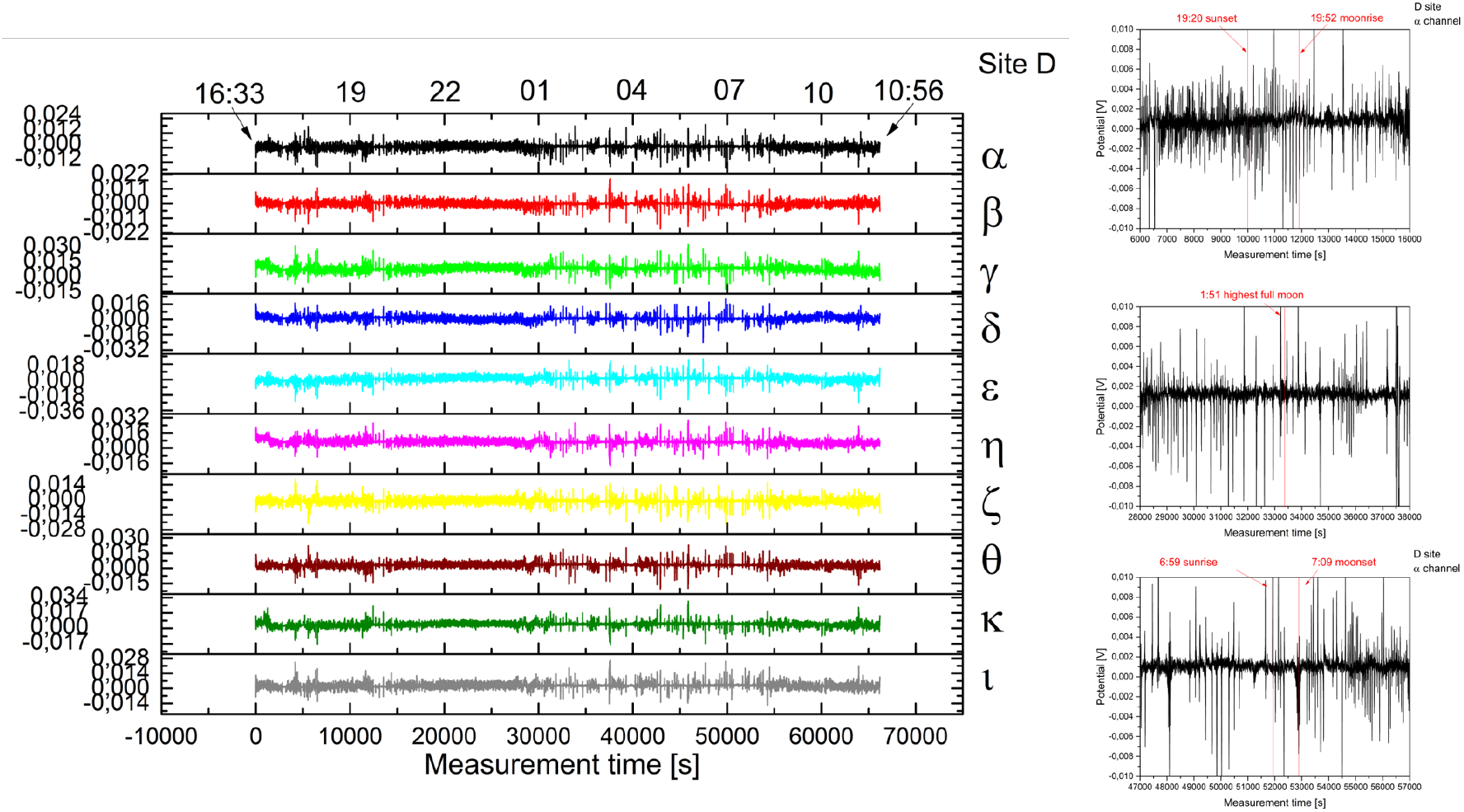
Data measured from site D, tree D1, over ten individual electrodes during the autumn equinox, with full moon and clear sky.

Channels show the following features (see Fig. 5): number of spikes, barcode entropy, Simpson index, Space filling, Kolmogorov entropy have the same features, while Signal entropy and Expressiveness have a structure that resembles to the modulus of the first derivative of the previous quantities (a peak can be observed where an increasing / decreasing slope is present); Tsallis and Rényi entropies appear to be poorly or not correlated to the previous entities. The number of spike-like peaks detected by the numerical algorithm is higher for channels closer to the leaves and minimal close to the roots; it shows the following structure: a basal line of number of spikes that has a maximum around midday, and a huge peak on top of the baseline with the maximum at midnight. This is less evident for channels close to the roots.

**Figure 5.**
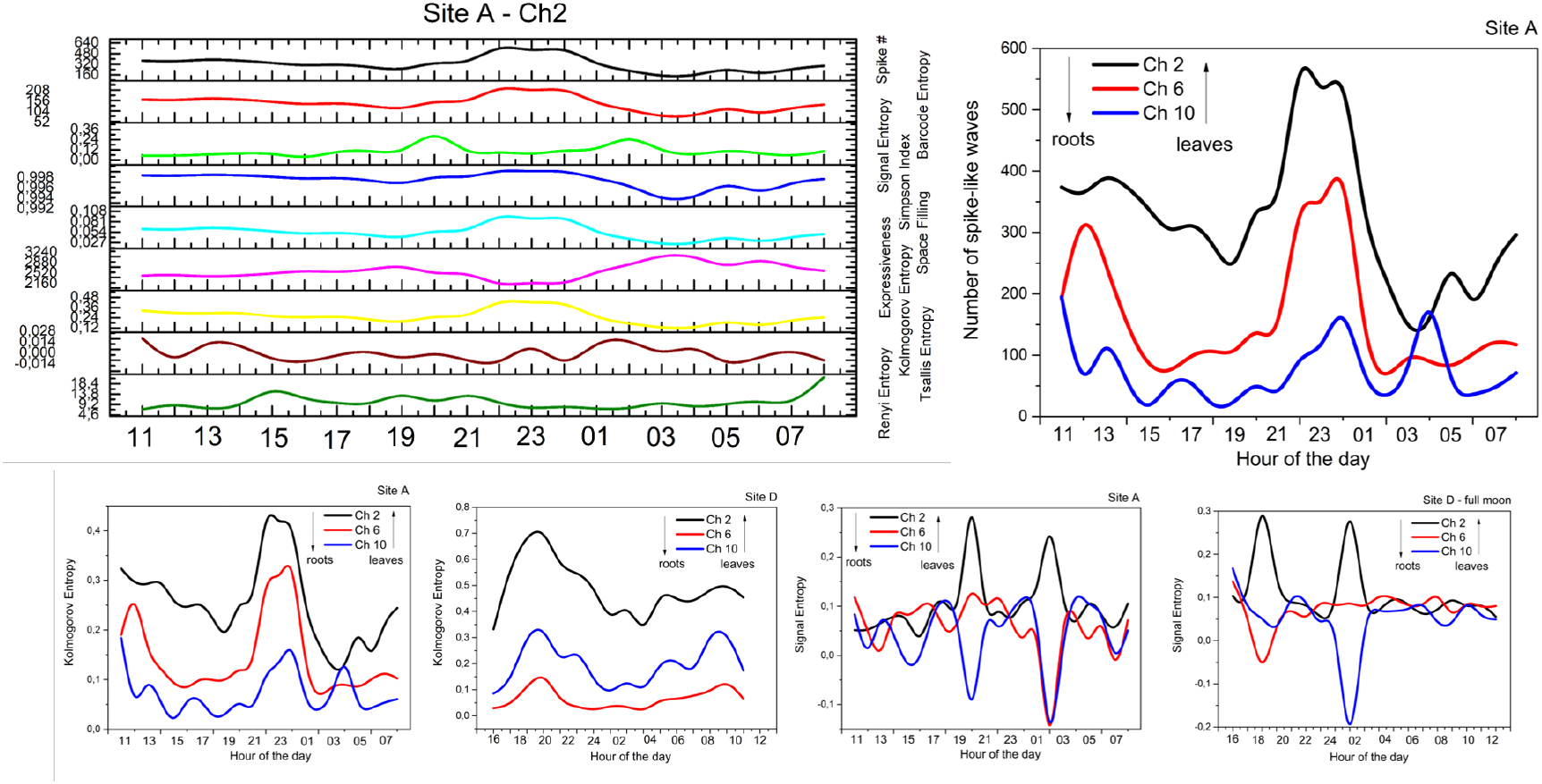
Top left: number of spikes, barcode entropy, Simpson index, space filling, Kolmogorov entropy, signal entropy, expressiveness, Tsallis entropy and Renyi entropy for a selected channel of a selected tree of a selected site. Top right: number of spikes for a selected channel of a selected tree of a selected site. Bottom, from left to right: Kolmogorov entropy of two selected channels of a selected tree of a selected site, Signal entropy of two selected channels of a selected tree of a selected site.

The signal entropy shows a correlation among all channels between the same site, though it is not possible to conclude anything relevant from the curve analysis. Kolmogorov entropy shows a structure similar to the number of spikes, featuring slightly less fluctuations and a clean curve that can well track the plant metabolic activity, or response to atmospheric phenomena. We can see that normally the channel closer to the leaves features higher entropies and that a structure with two peaks per day is preserved. Signal entropy shows very interesting features. Both site A and D have two peaks around 7 p.m. and 2 a.m.. Also site D shows a negative signal entropy for the log-positioned channels.

Visualising the histograms reporting peak duration and amplitudes one might discern between an active plant, featuring a higher diversity with both short spikes and longer periods, and a “dead” log, where the diversity is very much reduced (see Figure 6). The FFT shows a slow activity, particularly enhanced at frequencies around 100 mHz and below. A careful analysis of the E site, corresponding to a sector entirely occupied by dead logs, shows how some of them are locked and pretty much correlated. The two dimensional map proposed here shows the Kolmogorov entropy in colour scale, against measurement time and recording channel. Similar colour patterns against time are found in closer logs (# and * symbols).

**Figure 6.**
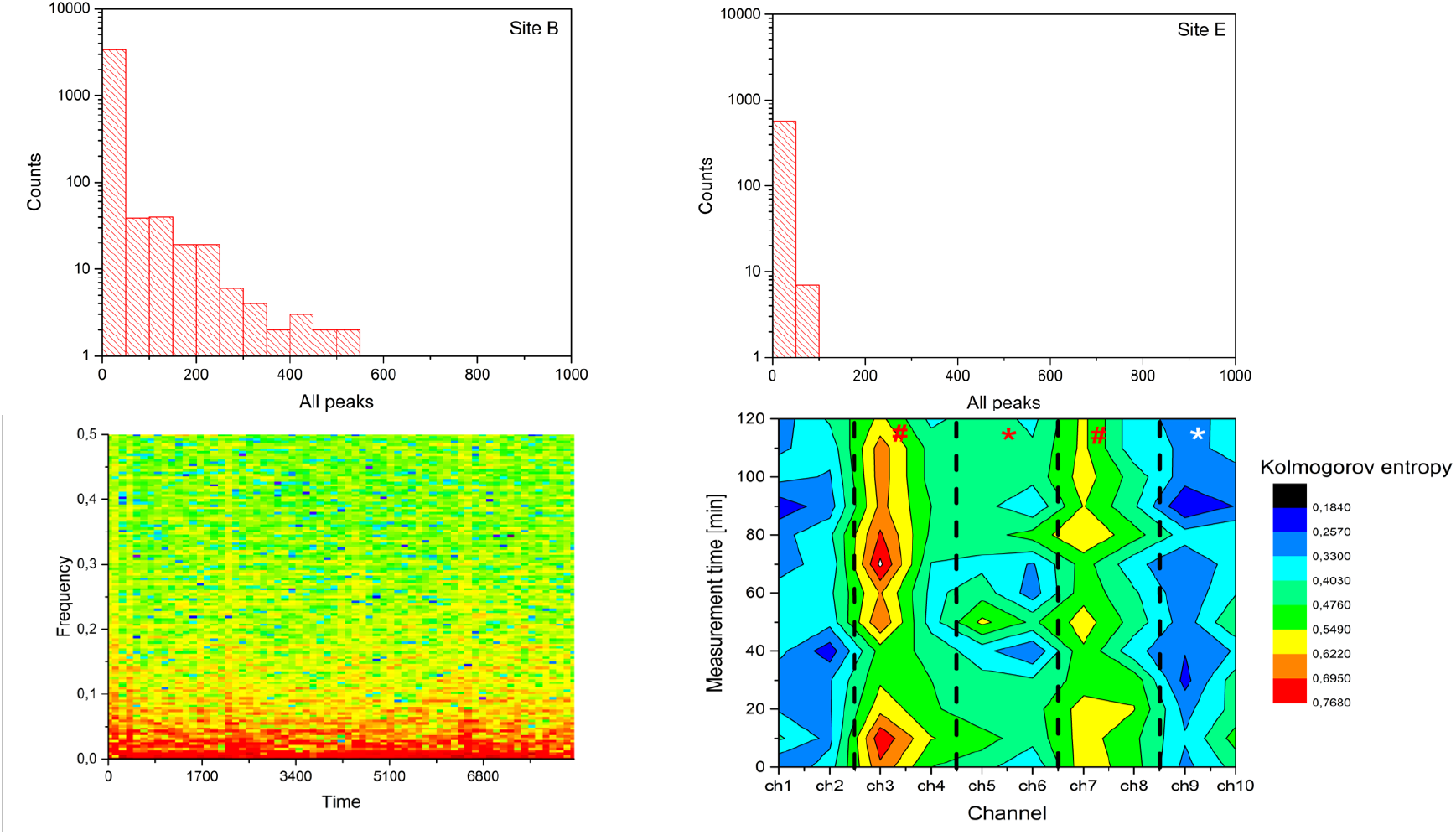
Top row: histograms of the population of spike-like structures detected on different sites. Bottom row, left: FFT map of a selected channel. Right: 2-dimensional map of the Kolmogorov entropy over the site populated by dead logs. Symbols identify correlated logs, dashed lines separate the couple of electrodes that work together in differential mode.

Resin samples from three conifer species (*Picea abies, Cedrus deodara*, and *Larix decidua*) have been characterized for their electronic transport properties, to assess if and how the production of such material by effect of the tree, in the bore hole and consequently in direct contact with the stainless steel electrodes, would influence the measurements. All the resins are highly aromatic and feature a complex composition: the *Picea abies* one is opaque, sticky and semi-solid, the *Cedrus deodara* one is clear, crystalline and fragile, and the *Larix decidua* one is cloudy and fluid as honey. There composition has been studied here [24] and here [25]. The impedance spectrum shows a broad range of behaviours (see Figure 7), from capacitive to inductive to neutral. In particular it has been found that the *Cedrus deodara*, in the investigated range of frequencies, behaves as an inductor, having a negative reactance that goes to zero at higher frequencies, while the *Picea abies* resin has a capacitive behaviour, having a positive reactance that goes to zero with increasing frequencies. The *Larix decidua* resin has a slightly capacitive behaviour, featuring a reactance that is hundred-folds smaller than that of the other species. Looking at the resistive component of impedance, we see that *Picea abies* and *Cedrus deodara* show similar values, while the *Larix decidua* sample is three to four orders of magnitude more conductive, probably owing to its liquid nature. Coming to the DC characterization, in the low voltage range applicable to the physiological potentials expressed by the trees, the resin is highly impeditive and shows a similar behaviour for all of the species, with some dispersion in the parameters extrapolated from linear fits to the experimental measurements: in particular for *Picea abies* the quality of the fit is reasonable (*R*^2^ = 0.8) and the slope (directly proportional to conductivity) is intermediate between the other two species. The high voltage behaviour also shows some interesting phenomenology, for what concerns *Picea abies* resin, where we can see a slight memristive behaviour, with a symmetrical, pinched hysteresis loop around 60 V of applied potential.

**Figure 7.**
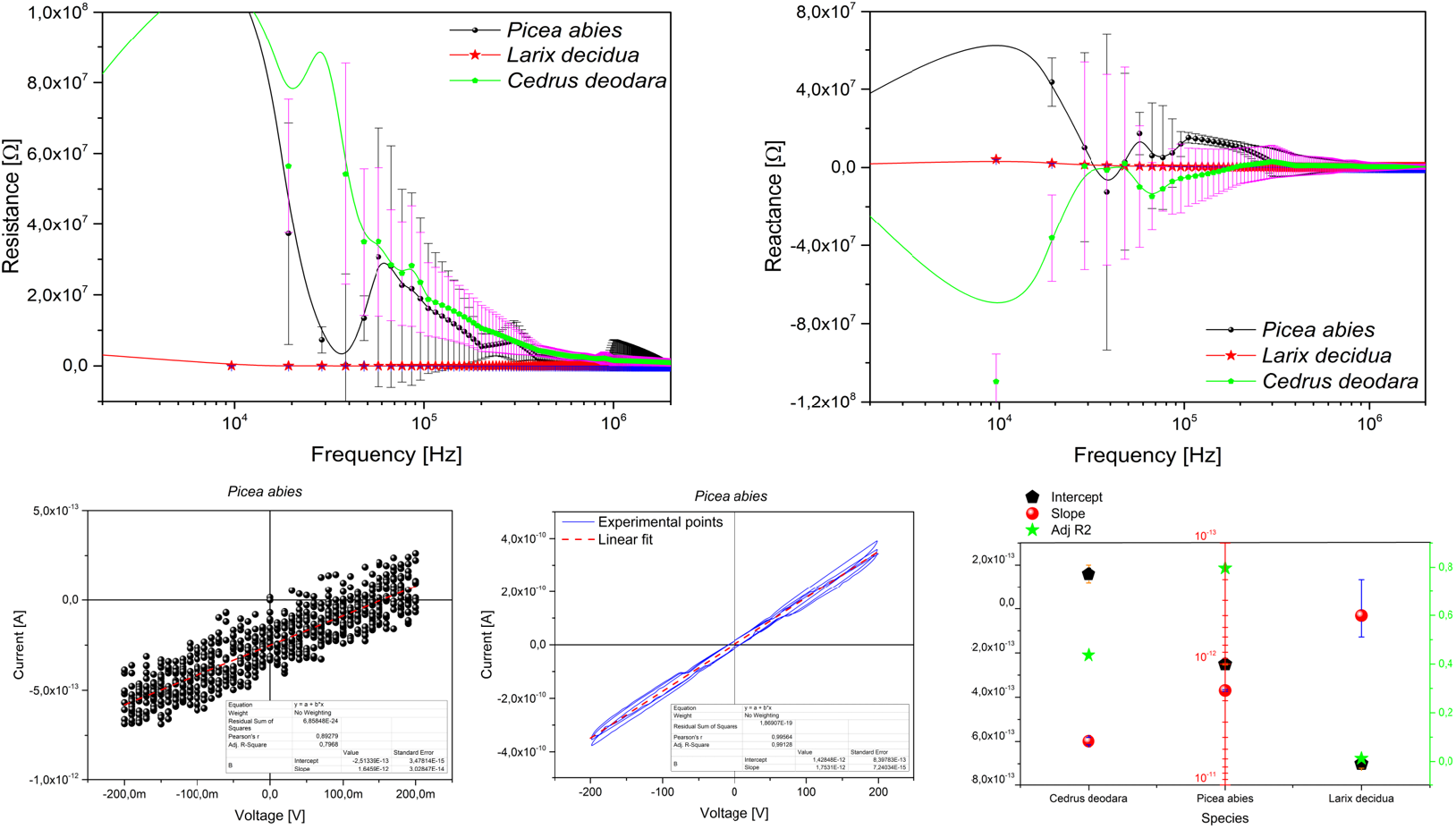
Electronic characterization of several resins: *Picea abies, Cedrus deodara*, and *Larix decidua*. Top row: impedance measurements, showing resistive (left) and reactive (right) components. Bottom row: low (left) and high (middle) voltage range behaviour of the *Picea abies* resin, and extrapolated linear fit parameters in a comparison between all the three resins (right).

## 4. An attempt to provide a theoretical modelling

### 4.1. Nambu-Goldstone fields and dissipative dynamics

Individual trees exchange energy and matter in different forms with their environment. They are in this sense “open” systems driven by dissipative dynamics under endogenous and external stimuli. We use the word ‘dissipative’ (and ‘dissipation’) in the sense that trees not only release to but also receive from the environment energy and matter.

In their dissipative dynamics, trees go through continuous adjustments of their equilibrium state, undergoing transitions through different dynamical regimes (*phase transitions*) triggered by changes in temperature, pressure, and other stimuli of external or endogenous origin. In each one of these regimes, the dynamical equilibrium with the environment is obtained according to general physical laws, as it happens in any other biological and physical system.

General laws of thermodynamics require the minimization of the free energy ℱ = *U* − *TS*, namely *d*ℱ = *dU* −*TdS* = 0, at each equilibrium (or quasi-equilibrium) state at a given temperature *T*, which expresses the first principle of thermodynamics for a system coupled to the environment at constant *T* and in the absence of mechanical work. For simplicity, we set equal to one the Boltzmann constant *k*_*B*_. *U* and *S* denote the internal energy and the entropy, respectively. Heat is *dQ* = *TdS*, as usual.

As well known, water plays a decisively central role in the tree vital processes and is subject to gradient forces of different origin (gravitation, piezometric pressure, heating and cooling gradients, chemical gradients, friction, etc.). The quantum electronic shells of water molecules present the charge distribution of an electric dipole. There is no preferred direction along which in the average the molecular dipoles point. The Lagrangian of the system, expressed in terms of these quantum dipole fields *ψ*(**x**, *t*), is thus invariant under the transformations of the SU(2) group of dipole spherical rotations. The polarization density *P* (**x**, *t*) of the system of water molecules is zero.

The dipole SU(2) symmetry is however *broken* by the mentioned forces acting on the water, which manifests in the non-zero *P* (**x**, *t*) of the system ground state, which therefore does not have the SU(2) symmetry of the Lagrangian [26, 27, 28, 29, 30].

The Breakdown mechanism of the Symmetry (BS) is well known in Quantum Field Theory (QFT). Here we do not consider the explicit BS induced by a change of the system Lagrangian, but the Spontaneous BS (SBS), where the symmetry breaking agent acts, as a trigger, not on the Lagrangian, but on the system states [31, 32, 33, 34].

According to the Goldstone theorem in QFT, SBS induces the dynamical formation of long range ordering correlations among the system components, i.e., in the present case, long range dipole waves among the molecular dipole fields.

The associated quanta are called Nambu-Goldstone (NG) fields, or quanta. In the QFT infinite volume limit, the NG quanta are massless boson modes that condense in coherent way in the system ground state. ‘Coherent’ means that long range correlations co-exist in the ordered patterns without negative interferences, so that the NG quanta, the Dipole-Wave-Quanta (DWQ) in the present case, behave approximately as an ideal gas of (quasi-)free particles. A measure of the ordering degree of the dipoles is provided by the non-zero value of a field, in the present case the polarization density *P* (**x**, *t*), called the ‘order parameter’.

Boundary effects (due to surfaces, impurities, etc.) may result in the non-zero effective mass *m*_*eff*_ of the DWQ fields. Dipole wave correlations extend accordingly over coherent domains of finite size of the order of *R* = *ħ /*(*m*_*eff*_ *c*), where *c* is the speed of light, *ħ* = *h/*(2*π*), and *h* the Planck constant.

The de Broglie wavelength of the DWQ quantum of momentum *p* is *λ* = *h/p*. In the DWQ ideal gas approximation, we have **p**^2^*/*(2*m*_*eff*_) = (3*/*2)*k*_*B*_*T*, with *k*_*B*_ the Boltzmann constant. Using *R* = *nλ/*2, with *n* an integer number, we obtain [30]

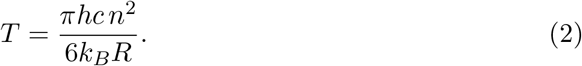

This equation shows that changes in temperature (due to dissipation) influence the range *R* spanned by dipole wave correlations, and viceversa; thus it sheds some light on the reported experimental observations of thermal spectra in connection with the trees reactivity to environmental stimuli.

Eq. (2) suggests that a control mechanism may be at work, aimed to keep constant the tree temperature, or to limit its variations within an interval convenient for the tree survival (also consistently with Le Chatelier principle on the system reactions opposing to external actions perturbing its (thermodynamic) equilibrium [35]). Suppose that an external agent produces an increase (decrease) in the temperature. Then the tree dynamical reaction aimed to keep *T* constant will *oppose* a decrease (increase) of *T* by increasing (decreasing) the size *R* of the coherent dipole wave propagation, according to Eq. (2). What happens at night, when external temperature drops, is counterfaced by plants decreasing *R*, which is then reflected in a higher frequency electrical activity, as discussed under Sect. **??**. It is clear, but worth to be stressed again, that in the described processes a crucial role is played by dissipativity, namely that the system is an open system, in permanent interaction with the environment.

We show that Eq. (2) expresses indeed, at the microscopic dynamical level, the free energy minimization constraint for the equilibrium state.

Consider variations of *T*, for given *n*, by differentiating Eq. (2):

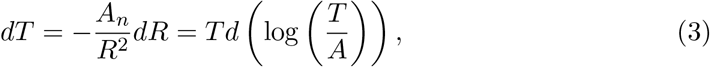

where it has been put *A*_*n*_ = *πhc n*^2^*/*(6*k*_*B*_) and Eq. (2) has been repeatedly used. Considering that *A*_*n*_, for given n, is constant, and since

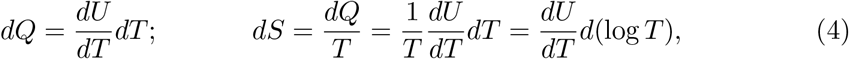

where partial derivative notation is not used since we are at constant volume *V*, Eq. (3) is rewritten as

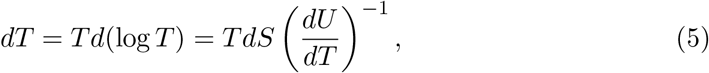

which is indeed the minimization condition of the free energy at constant *V*

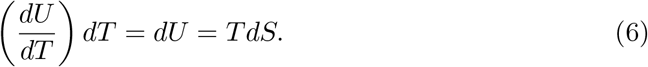

Note that the specific heat at constant *V* of our ‘ideal gas’ of dipole correlation quanta is of course *C*_*V*_ = *dQ/dT* = *dU/dT*. From Eqs. (3), (5) and (6) we see that variations in *T* actually imply entropy variations *dS*, namely ordering/disordering (*S* decrease/increase) through variations *dR* of the range over which dipole correlations extend. In these processes, there is energy ‘transmutation’ *U* ↔*TS*, of part of the energy from configurations with higher internal (kinetic) energy content, to more ‘ordered’ ones or, viceversa, part of the energy ‘stored’ in the ordering correlations is ‘disinvested’, released to obtain higher kinematic freedom (shorter range or loss of correlations, *S* increases).

We thus see that a remarkable dynamical internal degree of freedom is built in the system. The single equation *d*ℱ = 0 for the equilibrium constraint at constant *T* and *V*, is of course not enough to uniquely determine the values of the two variables *dU* and *dS*. Thus, the system dissipative dynamics, namely its being ‘open’ to external inputs, allows that a transfer (transmutation) of energy is allowed from random kinematics to order and viceversa, as described by ‘moving’ on the straight line of slope *T* = *dU/dS* in the plane (*U, S*).

The reported experimental measurements of Fig. 8 show that the response by the trees to the environment’s large gradients of temperature is such that their temperature does not reach the environment temperature. Instead, a sort of defense reaction or control mechanism is activated thus avoiding the tree’s excessive warming up or cooling down, possibly dangerous to its survival. According to the needs (dissipation), energy is stored in the ordering coherent dipole wave correlations, or taken from them.

**Figure 8.**
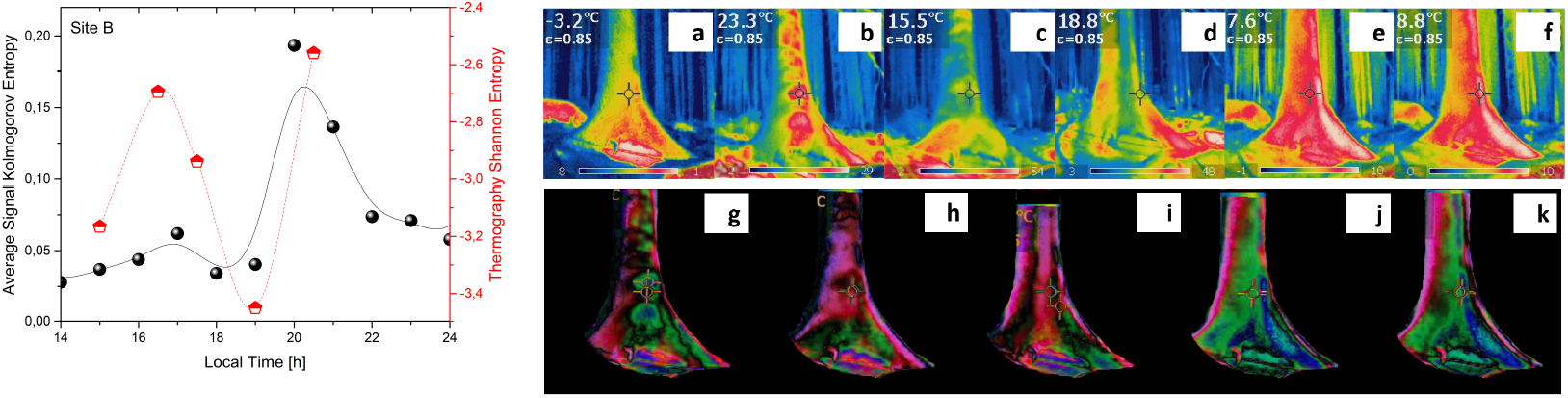
Left: qualitative matching between Kolmogorov entropy computed on the biopotential signal, averaged over the electrodes of site B, and Shannon entropy computed on the thermographies shown in the right panel, shooted from site B. Right: thermographies of a *Picea abies* trunk approximately 100 years old in site B (a to f) and algebraic difference between the registered frames above, to highlight changes in temperature distribution (g to k).

The entropic fingerprint of thermographies, if compared to that of the electrical signals recorded (see Fig. 8), shows quantitatively the same features, highlighting the metabolic and connection activity of trees.

### 4.2. Electromagnetic field and collective dynamical effects

In order to study the connection with the reported measures of the electromagnetic (em) field, we limit our discussion to the space components of the em vector potential **A**(**x**, *t*) of *A*_*µ*_(**x**, *t*) and use the Coulomb gauge condition **∇ · A**(**x**, *t*) = 0. When the em field is considered in the Lagrangian, this is invariant under local (and global) U(1) gauge transformations

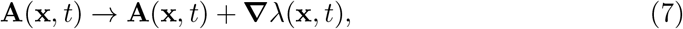

with **∇**^2^*λ*(**x**, *t*) = 0 imposed by the condition **∇ · A**(**x**, *t*) = 0. The global U(1) symmetry of the ground state is the one surviving the SBS of the spherical SU(2) symmetry of the dipole field *ψ*(**x**, *t*). However, also this global U(1) symmetry is spontaneously broken since it would be not possible to change simultaneously at every space-point by a constant amount *λ*, the phase of the dipole field *ψ*(**x**, *t*) in the ground state. SBS of the global U(1) is expressed by the non-zero value of the order parameter *v*(**x**, *t*), related to *P* (**x**, *t*) as |*v*(**x**, *t*) | ^2^ = 2*P* (**x**, *t*) [30, 34].

The wave function *σ*(**x**, *t*) of the charge density *ρ*(**x**, *t*), is

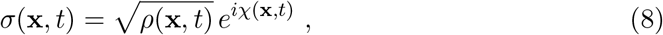

where the phase is the real field *χ*(**x**, *t*), and is related to *v*(**x**, *t*): |*σ*(**x**, *t*) |^2^ = *ρ*(**x**, *t*) ∝|*v*(**x**, *t*) |^2^. The phase *χ*(**x**, *t*) represents the NG wave field associated to the U(1) SBS [28, 30, 33]. The coherent condensation of the *χ*(**x**, *t*) quanta in the ground state is induced by the transformation

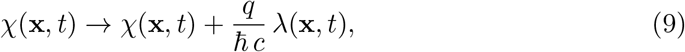

generating the (phase) transformation: *σ*(**x**, *t*) → exp[*i*(*q/ħ c*) *λ*(**x**, *t*)]*σ*(**x**, *t*).

The local gauge invariance of the Lagrangian allows the transformation **A**(**x**, *t*) → **A**^′^(**x**, *t*) = **A**(**x**, *t*)+(*ħ c/q*) **∇** *χ*(**x**, *t*), and then the transformation **A**^′^ (**x**, *t*) **A**^′^ → (**x**, *t*)+ **∇***λ*(**x**, *t*) is induced by the *χ*(**x**, *t*) boson condensation (9), with the constraint **∇**^2^*χ*(**x**, *t*) = 0 (and **∇**^2^*λ*(**x**, *t*) = 0) due to the gauge condition **∇ · A**(**x**, *t*) = 0 (**∇ · A**^′^(**x**, *t*) = 0).

Provided that such a gauge constraint is satisfied, *λ*(**x**, *t*) can be a constant or a space-time dependent function, producing homogeneous or non-homogeneous condensate structures in the system ground state, respectively.

For topologically non-trivial *λ*(**x**, *t*), boson condensation may describe vortices, rings and other extended objects with topological singularities [28, 32, 34, 36]. In these cases, non-commutativity of derivatives, e.g., (*∂*_1_ *∂*_2_ − *∂*_2_ *∂*_1_)*λ*(**x**, *t*) ≡ [*∂*_1_, *∂*_2_]*λ*(**x**, *t*) ≠ 0, denotes the non-equivalence between different paths connecting two points A and B and the em field *F*_*µν*_ is not invariant under the gauge transformation. Then, observable effects appear [28, 36]. On the contrary, no observable effects are produced by regular (non divergent or topologically trivial) gauge functions.

Since singularities are not compatible with non-zero mass *m*_*eff*_ of NG quanta, appearing due to boundary effects [32, 34, 36], topologically non-trivial extended objects cannot form near the system boundaries.

In the dissipative dynamics, mathematical consistency requires [32, 33, 34, 37] that the state of the system of NG dwq at temperature *T* is given by the two modes SU(1,1) generalized coherent state [38]

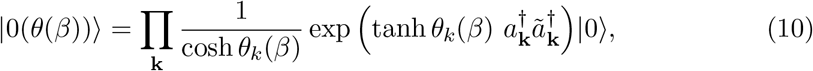

with *β* = 1*/k*_*B*_*T*, whose time dependence, *β* = *β*(*t*), is not explicitly shown for notational simplicity. |0(*θ*(*β*))⟩ is normalized to 1, ⟨ 0(*θ*(*β*)) | 0(*θ*(*β*)) ⟩ = 1, ∀*θ*(*β*), ∀*β*, ∀*t*. The operators *a*^†^_**k**_ and *a*_**k**_ are the NG dwq creation and annihilation operators, respectively, in terms of which the NG correlation field *χ*(**x**, *t*) is expanded. The operators 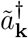 and *ã*_**k**_ are the creation and annihilation operators representing the tree’s environment (the reservoir or thermal bath). In QFT, finite temperature states are indeed condensates of the couples (*a*_**k**_, *ã*_**k**_), the”double” set of operators, those representing the system *a*_**k**_, and their “images” *ã*_**k**_, the “doubled” ones representing its environment. For formal details see [32, 33, 34, 37].

It can be shown that states at different temperatures, *T* |; *T* ^′^, are orthogonal states in the infinite volume limit *V* → ∞:

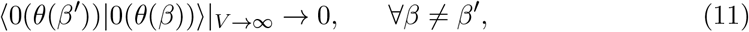

or, in the jargon of QFT, they belong to unitarily inequivalent representations of the canonical commutation relations (CCR), which means that they describe different dynamical regimes. This formally specifies the meaning of ‘different dynamical regimes’ referred to above. As said, at constant volume *V* and given *T*, the equilibrium constraint *dU* = *TdS* is described as the straight line of slope *T* in the (*U, T*) plane. The representation of the CCR at a given *β* (given *T*) describes the states on such a straight line. A change *β* →*β*^′^ leads to a different dynamical regime |0(*θ*(*β*))⟩ →|0(*θ*(*β*^′^)) ⟩, i.e. a change in the slope of the straight line in the (*U, T*) plane, a new ratio *dU/dS* between internal (kinetic) energy and energy stored in the dipole wave ordering correlations.

Entropy *S*_*a*_ and *S*_*ã*_ can be defined for each of the modes *a*_**k**_ and *ã*_**k**_, respectively, they have the same expression, with *ã*_**k**_ substituting *a*_**k**_, and *S*_*a*_ − *S*_*ã*_ is a conserved quantity. One finds:

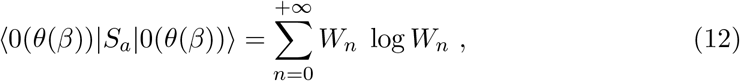

and similarly for *S*_*ã*_, with

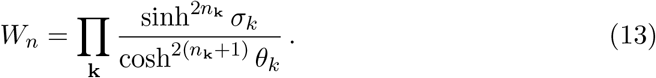

where *n* stays for the set {*n*_**k**_}, 0 *< W*_*n*_ *<* 1 and 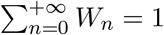 One also finds

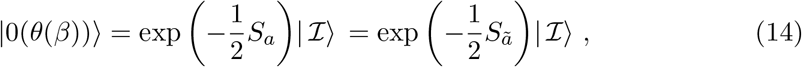

where 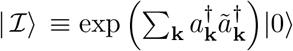, and

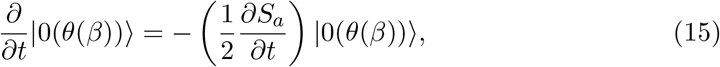

with *β* = *β*(*t*), and similar expression holds with *S*_*ã*_. Irreversibility of time evolution (*the arrow of time*) thus naturally emerges in the formalism. Eq. (15) shows indeed that entropy controls time evolution, consistently with the dissipative character of the dynamics. The free energy for the *a* modes is

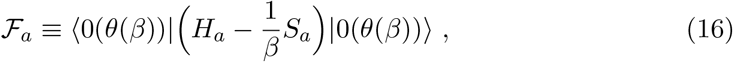

where 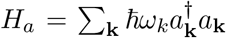, and similarly for *ã* modes. In the (quasi-)stationary case, the minimization condition *∂*ℱ_*a*_*/∂θ*_*k*_ = 0, ∀ *k*, leads to the Bose-Einstein distribution for *a*_**k**_ at time *t* in |0(*θ*(*β*))⟩:

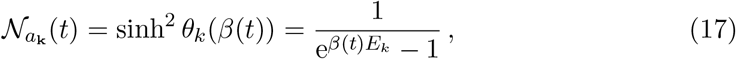

where *E*_*k*_ ≡ *ħω*_*k*_, and 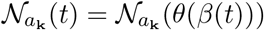, is the *a*_**k**_ number condensed in |0(*θ*(*β*))⟩. Also, *d*ℱ_*a*_ = *dE*_*a*_ − (1*/β*)*d*𝒮_*a*_ = 0 gives

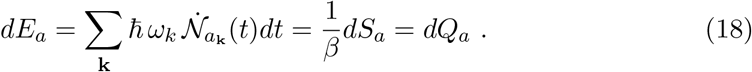

where 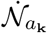 is the time derivative of 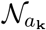. Eq. (18) explicitly shows that time variations *d*𝒩_*a*_ of the number of *a*_**k**_ modes condensed in the ground state |0(*θ*(*β*))⟩ manifest as heat *dQ*_*a*_ (and viceversa). These variations of 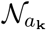 are actually variations in the dipole wave ordering correlations and produce changes by the same quantity (in sign and amount) *dQ*_*a*_ in internal energy *dE*_*a*_ and in entropy *dS*_*a*_.

Since, as said, *S*_*a*_ − *S*_*ã*_ is a constant of motion, *dS*_*a*_ = *dS*_*ã*_, i.e. *dQ*_*a*_ = *dQ*_*ã*_, explicitly showing the (dissipative) coupling with the environment. The state |0(*θ*(*β*))⟩ is in fact an entangled state for the *a*_**k**_ and *ã*_**k**_ modes. A quantitative measure of the entanglement is given by the linear correlation coefficient *J* (*N*_*a*_, *N*_*ã*_) [39, 40] (for simplicity we omit the subscript **k**):

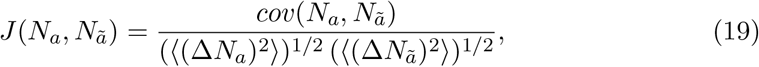

defined for non-zero values of ⟨ (Δ*N*_*a*_)^2^⟩ and ⟨ (Δ*N*_*ã*_)^2^⟩, with ⟨ (Δ*N*)^2^⟩ ≡ ⟨ (*N* − ⟨ *N* ⟩)^2^⟩ = ⟨ *N* ^2^⟩ − ⟨ *N* ⟩ ^2^ for the variance; the covariance is given by *cov*(*N*_*a*_, *N*_*ã*_) ≡ ⟨ *N*_*a*_*N*_*ã*_⟩ − ⟨ *N*_*a*_ ⟩ ⟨ *N*_*ã*_⟩. The symbol ⟨ ∗∗⟩ denotes expectation value in |0(*θ*(*β*)) ⟩. For non-correlated modes it is ⟨ *N*_*a*_*N*_*ã*_⟩ = ⟨ *N*_*a*_⟩ ⟨ *N*_*ã*_⟩ and the covariance is zero. For | 0(*θ*(*β*)) ⟩ it is *J* (*N*_*a*_, *N*_*ã*_) = 1. Recalling that *a*_**k**_ and *ã*_**k**_ are modes of the dwq field *χ*(**x**, *t*), which is a “phase” field (it appears in the phase, cf. Eq. (8)), we have shown that entanglement describes phase correlations.

The emerging picture is the one of the tree entangled, through its microscopic dynamics, with its environment, ‘in-phase’ with it. In the case other trees are in the environment, the entanglement “among trees” sheds new light on the ‘notion’ of *forest*, in some sense its ‘definition’, *the forest as an in-phase collective dynamical system*.

## 5. Conclusions and future prospects

We have shown the first outdoor installation measurements of patterns of electrical activity in a *Picea abies* forest, giving proofs of the following: the physiological activity of plants triggered by photosynthesis can be tracked, dead logs still kept alive by the nearby living plants can provide electrical signals correlated to their activity. Side characterizations show also the moon effect on the bio-electric potentials, and the electronic transport features of plant resin.

While many novel technologies are being developed to improve our daily lives, we are suggesting in this experiment to implement a technology that goes into a radically new direction: understanding the language of trees, eventually of fungi, that have created a universal living network perhaps running on a common language (see Figure 9). Messages conveyed along this network could tell us a lot about the health status of the forest, eventually also warn us about forthcoming disasters or ecological threats. We will ultimately explore the feasibility of a living reservoir computing system, with near zero energy requirements.

**Figure 9.**
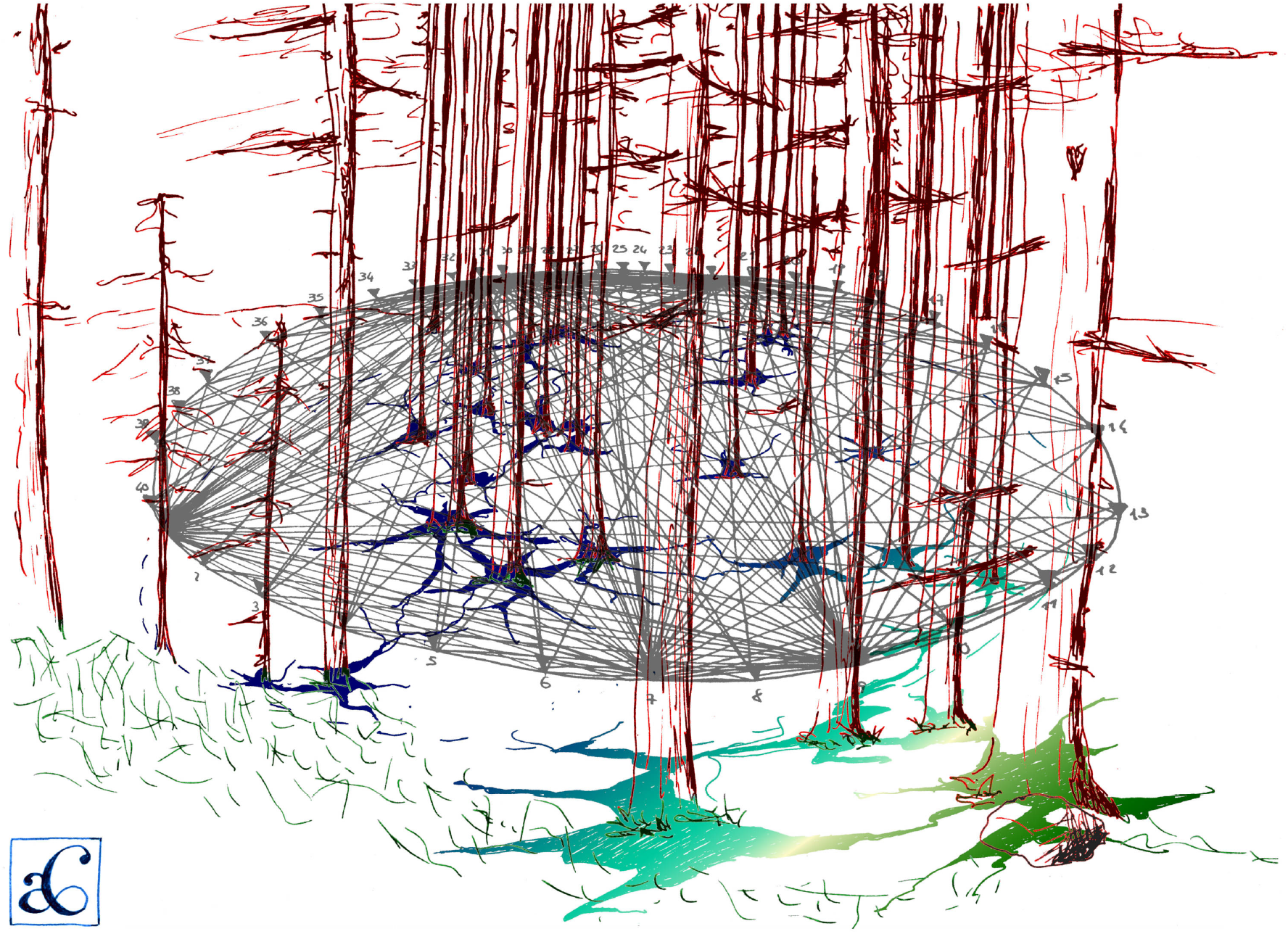
Artist view of the future measurements on the underground Wood Wide Web to map the network geometry by means of electrical tomography.

## Acknowledgements

The Authors wish to acknowledge the entire Corpo Forestale at Stazione Forestale Demaniale di Paneveggio e Cadino, in particular Girolamo Scarian, the entire workers squad, in particular Piero Baldessari, Dr. Andrea Daprà, Ente Parco Naturale di Paneveggio Pale di San Martino, Provincia Autonoma di Trento, the *Genius loci* and Salvanél, Magnifica Comunità della Valle di Fiemme, Prof. Renzo Motta and Prof. Francesca Secchi, Department of Agricultural, Forest and Food Sciences, University of Turin, Italy. A mention to our sponsors: OpenAzienda S.r.l.S., PrimoPrincipio Società Cooperativa, IGA Technology Services.

